# A genome-wide CRISPR screen identifies the glycosylation enzyme DPM1 as a modifier of DPAGT1 deficiency and ER stress

**DOI:** 10.1101/2021.12.03.471178

**Authors:** Hans M. Dalton, Raghuvir Viswanatha, Ricky Brathwaite, Jae Sophia Zuno, Stephanie E. Mohr, Norbert Perrimon, Clement Y. Chow

## Abstract

Partial loss-of-function mutations in glycosylation pathways underlie a set of rare diseases called Congenital Disorders of Glycosylation (CDGs). In particular, DPAGT1-CDG is caused by mutations in the gene encoding the first step in N-glycosylation, *DPAGT1*, and this disorder currently lacks effective therapies. To identify potential therapeutic targets for DPAGT1-CDG, we performed CRISPR knockout screens in *Drosophila* cells for genes associated with better survival and glycoprotein levels under DPAGT1 inhibition. We identified hundreds of candidate genes that may be of therapeutic benefit. Intriguingly, inhibition of the mannosyltransferase Dpm1, or its downstream glycosylation pathways, could rescue two *in vivo* models of DPAGT1 inhibition and ER stress, even though impairment of these pathways alone usually cause CDGs. While both *in vivo* models ostensibly cause ER stress (through *DPAGT1* inhibition or a misfolded protein), we found a novel difference in fructose metabolism that may indicate glycolysis as a modulator of DPAGT1-CDG. Our results provide new therapeutic targets for DPAGT1-CDG, include the unique finding of *Dpm1*-related pathways rescuing DPAGT1 inhibition, and reveal a novel interaction between fructose metabolism and ER stress.

## Introduction

Glycosylation comprises a variety of sugar-based post-translational modifications that occur primarily in the endoplasmic reticulum (ER) and Golgi apparatus^1^. It includes N-, O-, C-, and S-linked bonds as well as C-terminal-linked GPI anchors. Mutations in glycosylation pathways underlie rare diseases known as Congenital Disorders of Glycosylation (CDGs)^2^. CDGs can be caused by impaired enzyme activity or reduction of their specific metabolite substrates, such as N-Acetylglucosamine (GlcNAc), mannose, or uridine diphosphate (UDP). There are few treatment options available for these rare diseases, and research into the mechanisms of these pathways should help provide better avenues for advancing patient care.

Of the glycosylation pathways, N-linked glycosylation is critical for proper protein folding and function^1,3^. Inhibition of or abnormal N-linked glycosylation causes a build-up of misfolded proteins in the ER lumen, subsequent ER stress, and induction of the unfolded protein response (UPR)^3–8^. The UPR is a transcriptional response that attempts to restore homeostasis to the cell, but it can also lead to apoptosis when the stress is not resolved. The initial metabolite required for N-glycosylation is GlcNAc-PP-dolichol, which is synthesized by the dolichyl-phosphate N-acetylglucosamine phosphotransferase 1 (DPAGT1) enzyme^9^ (*Alg7* in *Drosophila*, hereafter referred to as *DPAGT1*). In humans, partial loss-of-function mutations in *DPAGT1* cause DPAGT1-CDG (CDG-Ij) with symptoms including seizures and developmental delay, among others^9,10^. Inhibiting DPAGT1 with the nucleoside Tunicamycin (Tun) reduces GlcNAc-PP-dolichol levels^11,12^ and induces ER stress^13^. DPAGT1-CDG patient cells, as well as *in vitro DPAGT1* mutant cell lines, are more sensitive to Tun^10,14^. *DPAGT1* mutant cell lines also show increased autophagy and signs of senescence^14^.

As with most CDGs, there are few therapeutic options available for treating DPAGT1-CDG. Any therapeutic option must be precise, as overexpression of DPAGT1 can also lead to improper N-glycosylation and may underlie some cancer phenotypes^15^. One alternative approach to current therapeutics is to identify modifier genes, where targeting genes that interact with *DPAGT1* expression or downstream phenotypes can potentially be used to develop new therapeutic options^16–18^. Moreover, determining if patients have differential baseline expression of these modifier genes may result in better personalized therapeutic solutions^17^.

Here we present genome-wide CRISPR screens for genes that modify phenotypes associated with reduced function of *DPAGT1*. These screens identify novel genes involved in modifying cellular and physiological outcomes associated with *DPAGT1* inhibition and increased ER stress. Of note, we find that knockout of multiple GPI anchor biosynthesis genes improves survival and cell surface glycoprotein levels associated with *DPAGT1* inhibition and ER stress. In addition, we find that the mannosyltransferase *Dpm1* is one of the strongest modifier genes and inhibition of *Dpm1* vastly improves cell survival under the loss of *DPAGT1* function and ER stress. We provide evidence that this effect likely occurs through the combined role of Dpm1 in O-mannosylation, N-glycosylation, and GPI anchor biosynthesis pathways. Importantly, the majority of our top candidate genes validate in *in vivo* models of *DPAGT1* inhibition and ER stress. In addition, results from the cell screens suggest that fructose metabolism might affect the loss of *DPAGT1* function, and we observed altered fructose metabolism in the *in vivo DPAGT1* model, suggesting that dietary fructose supplementation may hold therapeutic promise for these patients. Taken together, we identify numerous new candidate *DPAGT1* modifier genes that are potential therapeutic targets and report the remarkable finding that impairing CDG genes in parallel can result in an improved overall phenotype.

## Results

### Disruption of glycosylation and glycolytic pathways rescue the effects of *DPAGT1* inhibition

To identify genes impacting the loss of *DPAGT1* function and ER stress, we performed a cell-based, genome-wide CRISPR/Cas9 knockout screen (hereafter referred to as “survival screen”). Briefly, we used genome-wide single guide RNA (sgRNA) libraries transfected into a *Drosophila* S2 cell line to generate a pool of cells in which each cell carried gene knockouts (as described previously^19,20^, Fig. 1). Each gene tested had at least four different sgRNAs per gene in order to mitigate sgRNA-specific effects. After gene knockout, cells were either untreated or exposed to a sublethal dose of the DPAGT1 inhibitor Tunicamycin (Tun). After 30 days under DPAGT1 inhibition, cell populations were subjected to deep amplicon sequencing to determine sgRNA abundance. A change in the abundance of an sgRNA in the final population of DPAGT1-inhibited cells vs. untreated cells indicated that knockout of that gene caused resistance or sensitivity. Genes were ranked by the median log-fold change (lfc) in the abundance of each sgRNA targeting that gene (Table 1-2, Supp. Table 1).

**Table 1.** Summary of survival screen data.

| Resistance genes |  |  | Sensitivity genes |  |  |
| --- | --- | --- | --- | --- | --- |
| lfc | Total hits | % total genes | lfc | Total hits | % total genes |
| 2 or higher | 16 | 0.12% | -2 or lower | 1 | 0.01% |
| 1 – 1.99 | 156 | 1.18% | -1 – -1.99 | 9 | 0.07% |
| 0.5 – 0.99 | 979 | 7.40% | -0.5 – -0.99 | 241 | 1.82% |

**Figure 1.**
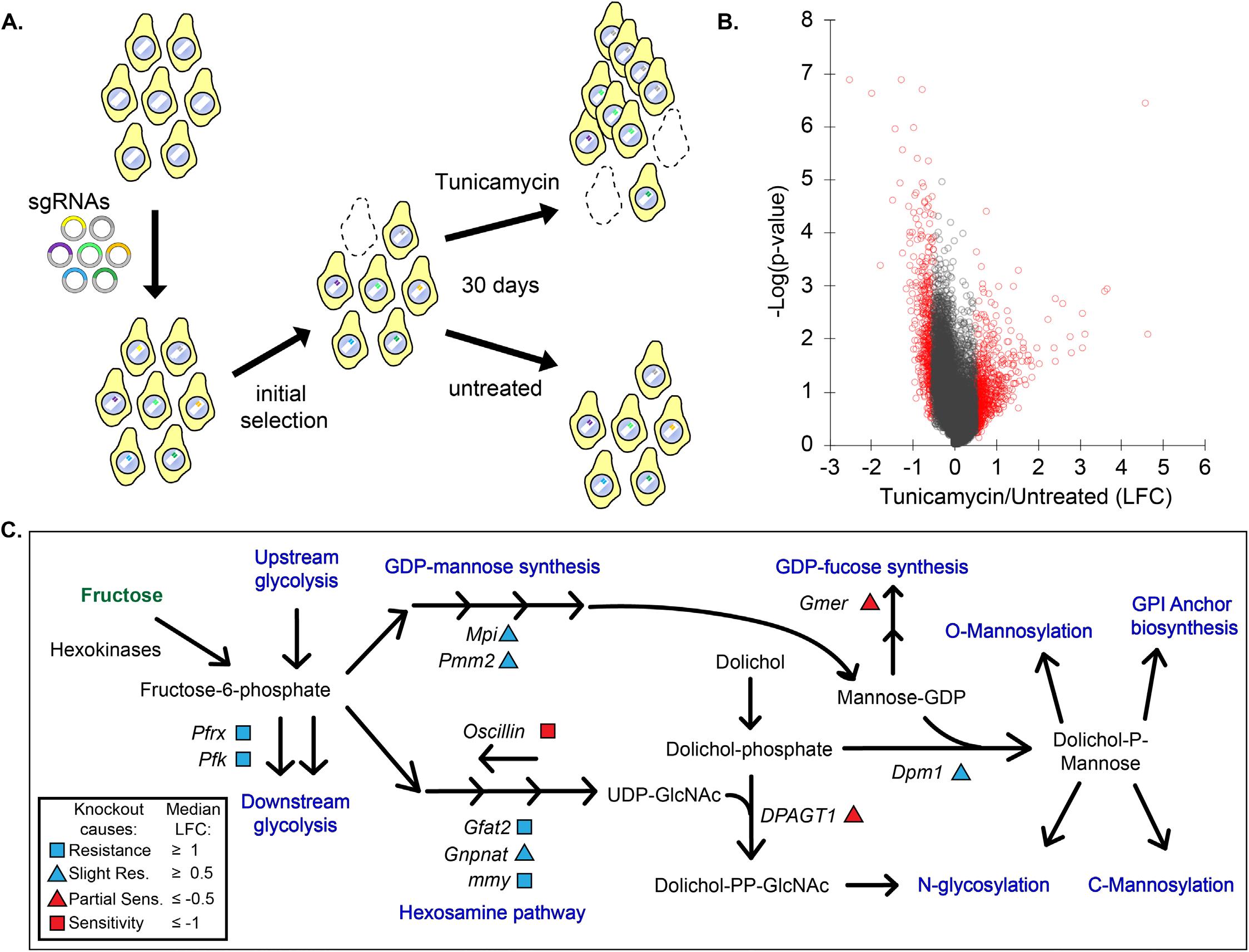
A survival screen reveals hexosamine, glycolysis, and GDP-mannose synthesis in resistance to *DPAGT1* loss. A. We introduced a whole genome guide RNA library into Drosophila cells stably expressing constitutive Cas9. Pooled cell populations were grown for 30 days either untreated or with Tun. Final cell populations were sequenced for sgRNA abundance to determine candidate genes implicated in Tun resistance or sensitivity. B. Volcano plot of survival screen. Red dots indicate genes with an absolute LFC value of at least 0.5. C. A simplified model of one set of highly enriched pathways for gene knockouts causing resistance or sensitivity. Note that only genes whose knockout provided resistance or sensitivity are labeled.

1151 gene knockouts (8.7% of genes tested) increased resistance to DPAGT1 inhibition at an lfc value of ≥0.5, while 251 gene knockouts (1.9% of genes tested) increased sensitivity to DPAGT1 inhibition at an lfc value of ≤-0.5 (Table 1). Top resistance genes include the major ER stress sensor *PEK* (human: *PERK;* lfc = 4.76), hexosamine enzymes *Gfat2* (*GFPT2*; lfc = 4.68) and *mmy* (*UAP1*; lfc = 1.57), the insulin-degrading enzyme *Ide* (*IDE*; lfc = 3.25), and the two glycolytic kinases *Pfk* (*PFKM*; lfc = 1.47) and *Pfrx* (*PFKFB1*; lfc = 1.74) (Table 2, Supp. Table 1). Top sensitivity genes include the proto-oncogene *pnt* (*ETS1*; lfc = −2.41), the hexosamine enzyme *Oscillin* (*GNPDA2*; lfc = −1.87), and the transporter-related genes *Rab6* (*RAB6A*; lfc = −0.85) and *Rab40* (*RAB40C*; lfc = −1.12) (Table 2, Supp. Table 1).

**Table 2.** Top 20 candidate genes providing resistance and sensitivity. Human orthologs are based on the highest DIOPT score^21^.

| Fly Gene | Human ortholog | lfc |
| --- | --- | --- |
| *PEK* | *EIF2AK3* | 5.51 |
| *Gfat2* | *GFPT2* | 4.68 |
| *CG9609* | *GTF3A* | 3.78 |
| *Hrd3* | *SEL1L* | 3.71 |
| *Ide* | *IDE* | 4.52 |
| *drongo* | *AGFG1* | 3.19 |
| *CG30345* | *SLC46A1* | 3.17 |
| *EcR* | *NR1H3* | 2.94 |
| *ham* | *MECOM* | 2.91 |
| *srp* | *GATA1* | 2.72 |
| *sau* | *GOLPH3* | 2.53 |
| *Pten* | *PTEN* | 3.23 |
| *Sin3A* | *SIN3A* | 2.52 |
| *Cdep* | *FARP2* | 2.37 |
| *CG10960* | *SLC2A8* | 2.16 |
| *REG* | *PSME3* | 2.08 |
| *dl* | *REL* | 1.98 |
| *ND-MLRQ* | *NDUFA4* | 1.97 |
| *CG2926* | *PHRF1* | 1.94 |
| *Gpdh1* | *GPD1* | 1.90 |
| *pnt* | *ETS1* | -2.41 |
| *Oscillin* | *GNPDA2* | -1.87 |
| *upSET* | *KMT2E* | -1.66 |
| *CG9590* | *FAM114A2* | -1.37 |
| *GalT1* | *B3GALT1* | -1.31 |
| *Asx* | *ASXL3* | -1.18 |
| *CG10984* | *ANKRD12* | -1.17 |
| *Vap-33A* | *VAPB* | -1.12 |
| *Rab40* | *RAB40C* | -1.12 |
| *Cdk2* | *CDK2* | -1.05 |
| *bel* | *DDX3X* | -0.98 |
| *P5cr-2* | *PYCR3* | -0.96 |
| *Tango14* | *NUS1* | -0.95 |
| *vtd* | *RAD21* | -0.95 |
| *CG33110* | *ELOVL1* | -0.95 |
| *CG32079* | *SLC36A2* | -0.95 |
| *Tor* | *MTOR* | -0.94 |
| *Oatp58Da* | *SLCO1A2* | -0.93 |
| *dre4* | *SUPT16H* | -0.90 |
| *sws* | *PNPLA7* | -0.90 |

Many top candidate genes are *Drosophila* orthologs of known CDG genes, such as *Gfat2*. To explore this further, we queried all known *Drosophila* CDG orthologs and a CDG genetic panel^22,23^ (145 genes, Supp. Table 2). We considered genes that increased sensitivity separately from genes that increased resistance. There was no significant enrichment of CDG genes among genes that increased sensitivity (6 observed vs. 2.7 expected, p=0.06, Supp. Table 2). Remarkably, there was a significant enrichment of CDG genes among genes that increased resistance to *DPAGT1* inhibition (23 observed vs. 12.6 expected, p<0.01, Supp. Table 2). This suggests that perturbation of certain CDG-related pathways in tandem can result in an overall improvement to cellular health.

We performed Gene Ontology (GO) analysis on all candidate genes with an absolute lfc ≥ 0.5 compared to untreated cells (Table 3, Supp Table 3). We chose to use both resistance and sensitivity genes in the same analysis as many pathways include genes that are either positive or negative regulators of the pathway. The top enriched categories included “nucleotide-sugar biosynthetic process” (GO:0009226) and “pyruvate metabolic process” (GO:0006090). These include genes involved in the hexosamine pathway and glycolysis, respectively, and relate to *DPAGT1*, as both of these pathways are critical in synthesizing upstream metabolites for several glycosylation pathways^24^. To further explore these connections, we examined an adjacent pathway (GO term “GDP-mannose metabolic process”, GO:0019673). This pathway involves the creation of GDP-mannose from fructose-6-phosphate, which involves the most common CDG gene *Pmm2* (PMM2; lfc = 0.50) and is also important in glycosylation pathways. Here, we found multiple genes at an lfc value cutoff of ±0.5 (4/7 *Drosophila* orthologs, Fig. 1C). Overall, there is strong enrichment of genes in glycolytic, hexosamine, and related pathways, suggesting the hypothesis that suppression of these pathways rescues the effects of *DPAGT1* inhibition.

**Table 3.** Gene ontology analysis of top candidate genes from the survival screen. Candidate genes with an lfc of ≥0.5 or ≤-0.5 were analyzed together.

| Gene Ontology | Fold Enriched | FDR |
| --- | --- | --- |
| nucleotide-sugar biosynthetic process (GO:0009226) | 6.57 | 3.85E-02 |
| nucleotide-sugar metabolic process (GO:0009225) | 6.34 | 5.26E-03 |
| regulation of feeding behavior (GO:0060259) | 4.38 | 4.40E-02 |
| female germ-line stem cell population maintenance (GO:0036099) | 3.94 | 1.11E-02 |
| larval midgut histolysis (GO:0035069) | 3.52 | 4.55E-02 |
| cuticle pattern formation (GO:0035017) | 3.39 | 3.74E-02 |
| midgut development (GO:0007494) | 3.38 | 2.58E-02 |
| pyruvate metabolic process (GO:0006090) | 3.37 | 1.78E-02 |
| cell surface receptor signaling pathway involved in cell-cell signaling (GO:1905114) | 3.29 | 4.42E-02 |
| synaptic assembly at neuromuscular junction (GO:0051124) | 3.29 | 4.40E-02 |

### Loss of GPI anchor biosynthesis rescues *DPAGT1* inhibition-induced reduction in cell surface glycoprotein levels

Tun inhibition of *DPAGT1* results in loss of cell surface glycoproteins^25,26^. Some candidate modifier genes might rescue the cellular survival phenotype by rescuing cell surface glycoprotein levels. To identify this subset of candidates, we performed a parallel genome-wide screen (Fig. 2). We used an identical pool of CRISPR knockout *Drosophila* cells as described above, either kept them untreated or exposed them to Tun, and then treated cells with fluorescently-labeled Concanavalin A (ConA), which binds cell surface glycoproteins^25,27^. Cells were then sorted based on fluorescence, with a higher fluorescent signal indicating more cell surface glycoproteins and vice versa. This assay allowed us to determine if a gene knockout can rescue DPAGT1 inhibition-induced loss of cell surface glycoprotein levels.

**Figure 2.**
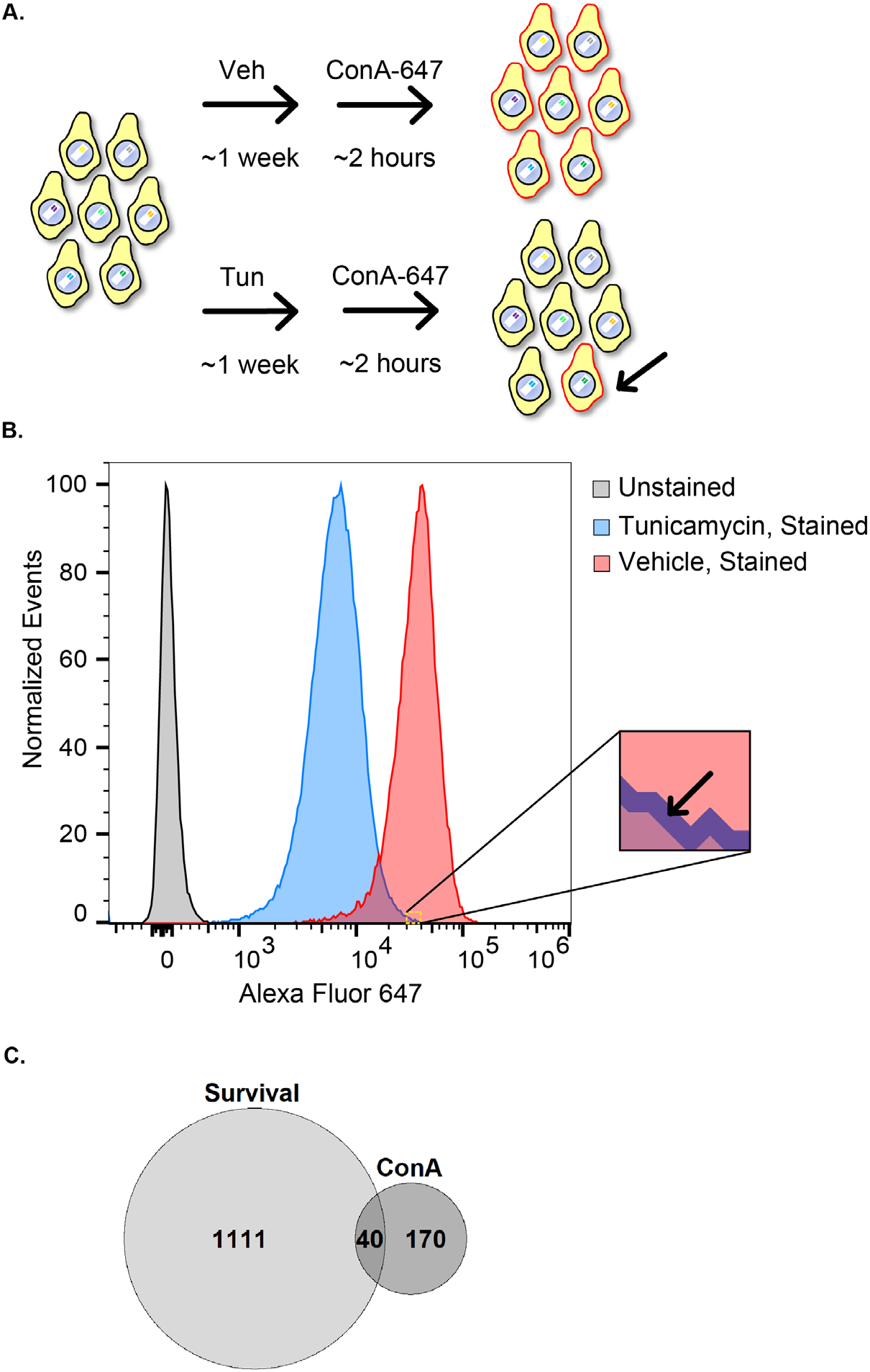
A Concanavalin A screen for gene knockouts capable of restoring cell surface glycoproteins under DPAGT1 inhibition. A. CRISPR knockout cells were either untreated or treated with Tun, and then stained with fluorescently-labeled ConA to mark cell-surface glycoproteins. B. Flow cytometry output of the ConA experiment. Tun treatment caused a ∼5-fold decrease in ConA staining intensity on average. The “blown-up” section denotes the start of the gate and encompasses Tun-treated cells capable of consistently restoring cell surface glycoproteins to untreated cell levels (the top ∼1% of cells). C. Venn diagram comparison of the survival and ConA screens. There were 40 genes that restored cell surface glycoproteins and provided resistance to DPAGT1 inhibition.

Under DPAGT1 inhibition, knockout of multiple genes was able to rescue cell surface glycoproteins to the levels observed in untreated cells (Supp. Table 4). We considered genes to be the strongest candidates if they had at least one sgRNA in the highest ∼1% of fluorescence of all sgRNAs in two biological replicates. (Fig. 2, Table 4, Supp. Table 4). This filtering resulted in identification of 209 genes (∼1.5% of all genes tested) whose knockout provided strong and consistent rescue of cell surface glycoprotein levels under DPAGT1 inhibition.

**Table 4.** Top 20 candidate genes from the ConA screen. Human orthologs are based on the highest DIOPT score^21^.

| **Fly Gene** | **Human ortholog** | **# Guides in top ~1%** |
| --- | --- | --- |
| *twin* | *CNOT6L* | 10 |
| *CG5276* | *CANT1* | 7 |
| *PIG-S* | *PIGS* | 6 |
| *PIG-A* | *PIGA* | 5 |
| *e(y)3* | *PHF10* | 5 |
| *HGTX* | *NKX6-1* | 5 |
| *slif* | *SLC7A1* | 5 |
| *ssh* | *SSH2* | 5 |
| *Afti* | *AFTPH* | 4 |
| *Bx* | *LMO1* | 4 |
| *CG17140* | *VDAC1* | 4 |
| *Mvk* | *MVK* | 4 |
| *dgt1* | *KANSL2* | 4 |
| *kay* | *FOS* | 4 |
| *mbt* | *PAK4* | 4 |
| *Nrg* | *NRCAM* | 4 |
| *par-6* | *PARD6G* | 4 |
| *PEK* | *EIF2AK3* | 4 |
| *Skp2* | *SKP2* | 4 |
| *uzip* | *None* | 4 |

GO analysis of the 209 genes that restore cell surface glycoprotein levels indicated that “GPI anchor biosynthetic process” (GO:0006506), “glycolipid metabolic process” (GO:0006664), and “inositol metabolic process” (GO:0006020) were the top enriched categories (Table 5, Supp. Table 5). Glycosylphosphatidylinositol (GPI) anchor biosynthesis is a form of glycosylation in which a GPI glycolipid is post-translationally attached to proteins in the ER. The first step in GPI anchor biosynthesis involves the attachment of GlcNAc to phosphatidylinositol, and the final GPI-associated proteins are typically transported to membrane-bound lipid rafts^28,29^. Phosphatidylinositol glycan (“*PIG*”) genes encode a diverse class of enzymes that build the GPI sugar chain^28,29^. Five different *PIG* genes were among the 209 hits, and all had at least 3 sgRNAs combined from both replicates, placing them in the top ∼22% of candidate genes. One of these is *PIG-A* (*PIGA)*, which was also a hit in the survival screen (lfc = 0.81). The other four were *PIG-S* (*PIGS), PIG-B* (*PIGB), PIG-H* (*PIGH)*, and *PIG-O* (*PIGO)*. Notably, these five *PIG* genes are spread throughout the entire GPI anchor biosynthesis process: PIGH and PIGA are both components of the initial GPI-GlcNAc transferase complex that attaches GlcNAc to phosphatidylinositol. PIGB performs an intermediate mannosylation step. PIGO attaches an ethanolamine after PIGB. PIGS is part of the final GPI-transamidase complex which attaches the GPI anchor to a protein. Given their diverse functions, spanning two separate enzyme complexes, it is likely that GPI anchor biosynthesis as a whole can influence the phenotype. Remarkably, the data suggest that impairment of GPI anchor biosynthesis, a process that is typically known to create cell surface glycoproteins, can rescue cell surface glycoprotein levels under DPAGT1 inhibition.

**Table 5.** Gene ontology analysis of candidate genes from the ConA screen.

| Gene Ontology | Fold Enrichment | FDR |
| --- | --- | --- |
| GPI anchor metabolic process (GO:0006505) | 10.82 | 1.46E-02 |
| liposaccharide metabolic process (GO:1903509) | 8.65 | 1.34E-02 |
| glycolipid metabolic process (GO:0006664) | 8.65 | 1.41E-02 |
| glycolipid biosynthetic process (GO:0009247) | 8.52 | 3.74E-02 |
| membrane lipid metabolic process (GO:0006643) | 7.06 | 2.03E-03 |
| membrane lipid biosynthetic process (GO:0046467) | 6.9 | 7.82E-03 |
| phosphatidylinositol metabolic process (GO:0046488) | 6.9 | 8.53E-03 |
| lipoprotein metabolic process (GO:0042157) | 6.31 | 4.67E-02 |
| glycerophospholipid metabolic process (GO:0006650) | 6.07 | 4.54E-03 |
| glycerolipid metabolic process (GO:0046486) | 5.9 | 1.85E-03 |

We hypothesized that some genes rescue the cell survival phenotype (Fig. 1) by rescuing cell surface glycoprotein levels. To identify these genes, we compared the 209 genes that rescue cell surface glycoprotein levels to the 1151 genes that increase resistance to DPAGT1 inhibition from the survival screen. Strikingly, 40 genes (∼19.1% of ConA hits; Supp. Table 6) — including top resistance hits (lfc > 1.5) *PEK, Ide*, and *Pfrx*, as well as *PIG-A* — overlapped, which is higher than expected by chance (2.2 fold higher, p<0.0001). Thus, a substantial percentage of gene knockouts that restore cell surface glycoproteins also increase resistance. Nevertheless, as the majority of genes that cause increased resistance did not have this effect (1111 genes, 96.5%), the consistent rescue of cell surface glycoprotein levels is not required to increase resistance to DPAGT1 inhibition.

### Inhibiting the mannosyltransferase Dpm1 rescues DPAGT1 inhibition and ER stress *in vivo*

To test candidate genes *in vivo*, we created a DPAGT1 disease model in *Drosophila* that expresses *DPAGT1* RNAi in the eye - causing a small, degenerate eye phenotype (hereafter referred to as “*DPAGT1* model”; Fig. 3A). Using the *DPAGT1* model, we can determine the impact of knocking down candidate genes with RNAi by measuring changes in the eye phenotype both quantitatively by size and qualitatively by observing visible phenotypes such as necrosis. We hypothesized that RNAi knockdown of candidate genes from the previous cell screens that improved phenotypes under DPAGT1 inhibition would improve the eye phenotype in the *DPAGT1* model. We primarily focused on the top GO enrichment categories from the survival screen (Table 3) and tested multiple RNAi lines against the glycolytic, hexosamine, and GDP-mannose-related pathways. We used multiple RNAi lines against each gene when available and/or used RNAi against multiple genes in the same pathway throughout - see Supp. Table 7 for complete RNAi line information.

**Figure 3.**
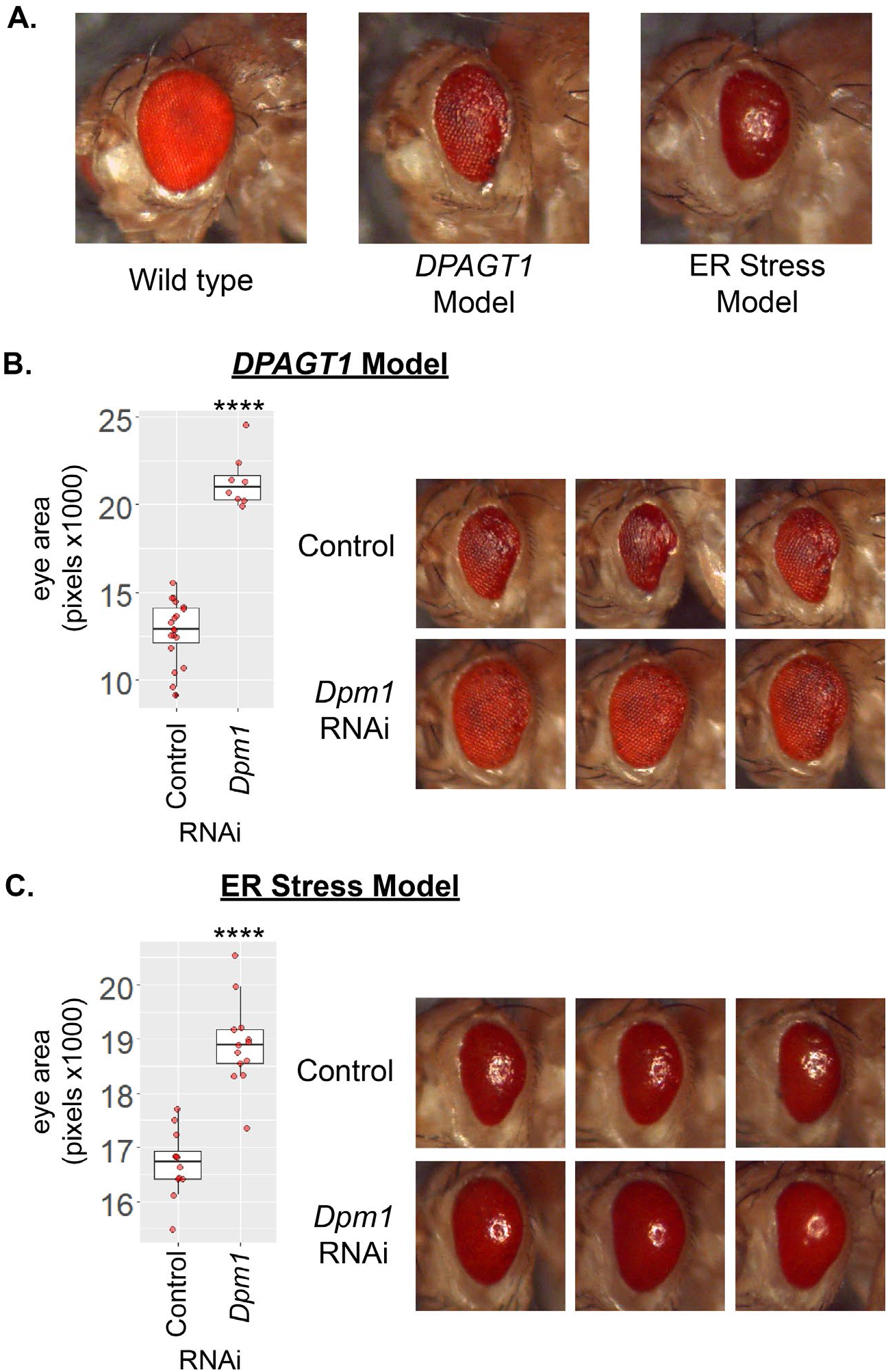
Knockdown of the mannosyltransferase *DPM1* rescues the *DPAGT1* and ER stress models. A. We generated a *DPAGT1* model that drives RNAi against *DPAGT1* in the fly eye (middle). This causes a degenerated eye phenotype. In addition to the *DPAGT1* model, we also used an ER stress fly model that overexpresses the misfolded protein Rh1^G69D^ in the eye (right). B. RNAi against the mannosyltransferase *Dpm1* strongly rescues the *DPAGT1* model both quantitatively and qualitatively. The three images are representative from the same experimental cross and RNAi line (BDSC 51396). **** p<0.0001 (Student’s t test). *C. Dpm1* RNAi rescues the ER stress model. Eye images are representative from the same experimental cross and RNAi line (BDSC 50713). **** p<0.0001 (Student’s t test).

We had initially tested 26 candidate genes *in vivo* (Supp. Table 8). Many of these candidate genes had *in vivo* effects that were consistent with the outcome from the cell screens, but others had no effect or even the opposite effect. However, some of the strongest candidate genes had an effect in the *DPAGT1* model that was consistent with the effects in the cell screens (Supp. Table 7). For example, we tested *Pfk* (lfc = 1.47) and *Pfrx* (lfc = 1.74), as they have the highest lfc values in the glycolytic pathway from the survival screen. *Pfrx* was also a hit in the ConA screen (Supp. Table 4). Matching the strong effect that we observed in the survival screen, RNAi against either of these two phosphofructokinases improved the *DPAGT1* model (*Pfk*: +32.2%, *Pfrx*: +11.3%, averaged across two biological replicates). On the other hand, in the hexosamine pathway, RNAi against transaminase-encoding *Gfat2*, despite causing resistance (lfc = 4.68), did not significantly affect the *DPAGT1* model (Supp. Table 7). Moreover, RNAi against the deaminase-encoding *Oscillin*, which caused sensitivity (lfc = −1.87), improved the *DPAGT1* model (+28.8%). These differences may be caused by the differing biology of a cell line vs an *in vivo* eye model.

RNAi against the mannosyltransferase *Dpm1* (*DPM1;* lfc = 0.68) showed the strongest rescue of the *DPAGT1* model phenotypes (+62.1%) (Fig.3B, Supp Table 7, Supp. Fig. 1-2). *Dpm1* encodes dolichol-phosphate mannosyltransferase 1, which, in complex with DPM2 and DPM3, synthesizes dolichol phosphate mannose (Dol-P-Man), an essential substrate in N-glycosylation, O-and C-mannosylation, and GPI anchor biosynthesis^30,31^ (Fig.1C). Loss of *DPM1* is usually detrimental to cells and patients^10,31,32^, yet knockdown of *Dpm1* improved the *DPAGT1* model.

To determine if the *Dpm1* rescue was specific to *DPAGT1* or ER stress in general, we also tested each RNAi line on the well-established *Drosophila* Rh1^G69D^ ER stress model^33–36^ (hereafter referred to as “ER stress model”) (Fig. 3A). The ER stress model overexpresses a misfolded protein in the eye in order to induce ER stress, apoptosis, and a degenerate eye phenotype. Similar to the *DPAGT1* model, *Dpm1* RNAi resulted in the strongest rescue of overall eye size in the ER stress model (+18%, Fig. 3C, Supp Table 7, Supp. Fig. 1-2), indicating that *Dpm1* inhibition is capable of rescuing the effects of *DPAGT1* inhibition and ER stress in general. Despite the fact that *Dpm1* is an essential enzyme, inhibiting *Dpm1* provides significant benefits to *DPAGT1* inhibition and ER stress, and thus, it may be a key factor in rescuing the effects of DPAGT1 deficiency.

### Impairment of O-mannosylation, N-glycosylation, and GPI anchor biosynthesis improves the loss of *DPAGT1* function and ER stress outcomes

One hypothesis for why inhibiting *Dpm1* might reverse *DPAGT1* inhibition and ER stress is that it would cause a reduction of its downstream product, Dol-P-Man, and in turn, impair downstream glycosylation pathways^37,38^. This is a particularly attractive hypothesis because, as shown above, under *DPAGT1* inhibition, loss of GPI anchor biosynthesis function, which utilizes Dol-P-Man in three separate steps, can improve cell survival and restore cell surface glycoproteins (Fig. 1-2, Supp. Table 1, 4). To test this hypothesis, we crossed strains expressing RNAi against the mannosyltransferase enzymes in O- and C-mannosylation and N-glycosylation to the *DPAGT1* model (Supp. Fig. 3-4). Because of the strong rescue effect of a number of GPI anchor biosynthesis genes, we also tested additional genes in this pathway (Supp. Fig. 3-4).

O-mannosylation begins with the addition of mannose to Ser/Thr residues, followed by a lengthening of the chain with additional sugar molecules^39,40^. One important O-mannosylated protein is alpha-dystroglycan, and impairment of O-mannosylation is associated with muscular dystrophy in humans^40^. The first step in O-mannosylation, which attaches the initial mannose from Dol-P-Man, requires an enzyme complex consisting of two O-mannosyltransferases, POMT1 and POMT2. RNAi against rotated abdomen (*rt, POMT1*) or twisted (*tw, POMT2*) rescued eye size in both the *DPAGT1* (*rt*: +6.2%; *tw*: +45.9%) and ER stress (*rt*: +10.2%; *tw*: +3.8%) models (Fig. 4A-B, Supp. Table 7, Supp. Fig. 3-4). These data suggest that loss of O-mannosylation can rescue the loss of *DPAGT1* function and ER stress.

**Figure 4.**
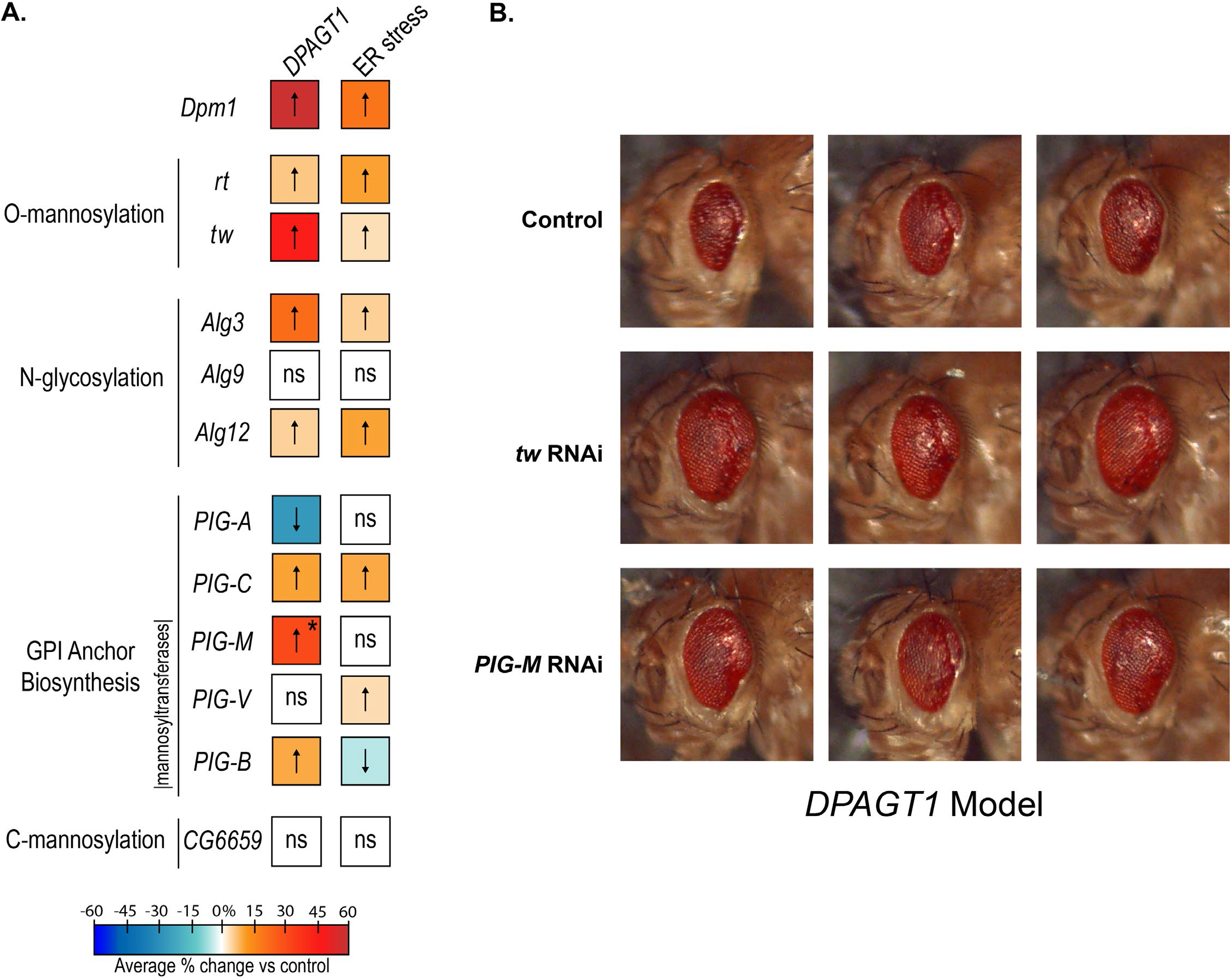
Knockdown of glycosylation pathways rescue the *DPAGT1* and ER stress models. A. A summary of RNAi data performed on downstream DPM1 glycosylation pathways. The majority of mannosyltransferases tested in O-mannosylation, N-glycosylation, and GPI anchor biosynthesis pathways increased eye size in the *DPAGT1* model. Colors denote % change of eye size vs their control; warmer colors (or up-arrows) have a stronger effect, and vice versa. Colors are derived from the averages of at least 2 biological replicates with at least one significant replicate and no opposite results (see Supp Table 7 for all RNAi data). * = One *PIG-M* RNAi line was very strong (BDSC 51890), while a second line was neutral/negative in response (see Supp Table 7). B. Representative images of the *DPAGT1* model crossed with RNAi against the two strongest hits, *tw* and *PIG-M* (shown are three representative images from the same RNAi line and experimental cross – *tw*: BDSC 55735, *PIG-M*: BDSC 51890)

In N-glycosylation, mannosylation steps utilizing Dol-P-Man occur in the ER lumen via the mannosyltransferases ALG3, ALG9, and ALG12^41^. RNAi against *Alg3* (*ALG3*) and *Alg12* (*ALG12*) improved eye size in both the *DPAGT1* (*Alg3*: +21.4%; *Alg12*: +4.5%) and ER stress (*Alg3*: +4.8%; *Alg12*: +9.9%) models. However, RNAi against *Alg9* (*ALG9)* did not affect either model (Fig. 4A, Supp. Table 7, Supp. Fig. 3-4). Overall, loss of the N-glycosylation mannosyltransferase steps can rescue the loss of *DPAGT1* function and ER stress.

In the GPI anchor biosynthesis pathway, most RNAi lines tested improved the *DPAGT1* model. These include a subunit of the GPI-GlcNAc transferase complex, *PIG-C* (*PIGC*, +22.6%), and the mannosyltransferases *PIG-B* (*PIGB*, +11.3%) and *PIG-M* (*PIGM*, +31.9%). A second *PIG-M* RNAi line had a smaller, negative effect (−5.7%). RNAi lines against the mannosyltransferase *PIG-V* (*PIGV*) did not have a significant effect. Surprisingly, RNAi against *PIG-A* was detrimental to the *DPAGT1* model (−5.7% and −27.7%, respectively). Unlike the previous two pathways tested, the ER stress model was not as consistent with the *DPAGT1* model, though it also had an overall beneficial effect (Fig. 4A-B, Supp. Table 7, Supp. Fig. 3-4). This may reflect a difference in how GPI anchor biosynthesis affects each model.

C-mannosylation is a less common type of glycosylation that involves the addition of mannose to tryptophan found in thrombospondin type 1 repeats^42^. The four enzymes in humans responsible for C-mannosylation are *DPY19L1-4* (*DPY19L1* and *DPY19L3* being the most functionally important^42^). In *Drosophila, CG6659* is the single ortholog of all four human paralogs (DIOPT scores for *DPY19L1-4*: 10/16, 9/16, 4/16 and 4/16). RNAi against *CG6659* did not affect eye size in either model; however, here we note an increased chance of a false negative as only one RNAi line and one gene from this pathway were examined (Fig. 4A, Supp. Table 7, Supp. Fig. 3-4). Thus, it is not immediately clear that impairing C-mannosylation imparts any benefit under the loss of *DPAGT1* function or ER stress.

Taken together, disruption of any of the three pathways downstream of Dpm1 and its product Dol-P-Man — O-mannosylation, N-glycosylation, and GPI anchor biosynthesis — can rescue the effects of *DPAGT1* inhibition and ER stress. Notably, RNAi against any single gene in these three pathways never reached the same magnitude of rescue provided by *Dpm1* RNAi (+62.1%) - with the closest being *tw* (+45.9%) and *PIG-M* (+31.9%) RNAi (Fig. 4B). This suggests that the strong rescue of the *DPAGT1* model with *Dpm1* RNAi may be from the synergy of the combined impairment of the three downstream glycosylation pathways. In addition, for every gene tested, RNAi downregulation in the *DPAGT1* model had stronger, more consistent rescue of negative phenotypes than the ER stress model, which suggests an underlying difference between the two models.

### Fructose metabolism is differentially altered depending on the source of ER stress

Functions related to glycolysis and metabolism were enriched among our candidate genes (Supp. Fig. 5, Table 3, Supp. Table 3). In the survival screen (Fig. 1), knockout of glycolytic genes generally caused resistance to *DPAGT1* inhibition, including the aforementioned phosphofructokinases *Pfk* (lfc = 1.47) and *Pfrx* (lfc = 1.74), as well as the dehydrogenase *Gapdh1* (*GAPDH*; lfc = 1.14), phosphoglycerate mutase *Pglym87* (*PGAM2*; lfc = 1.21), and pyruvate kinase *CG12229* (*PKM;* lfc = 0.73), among others (Supp. Table 1, Supp. Fig. 5). Thus, impairing glycolysis appears to improve survival under the loss of *DPAGT1* function.

**Figure 5.**
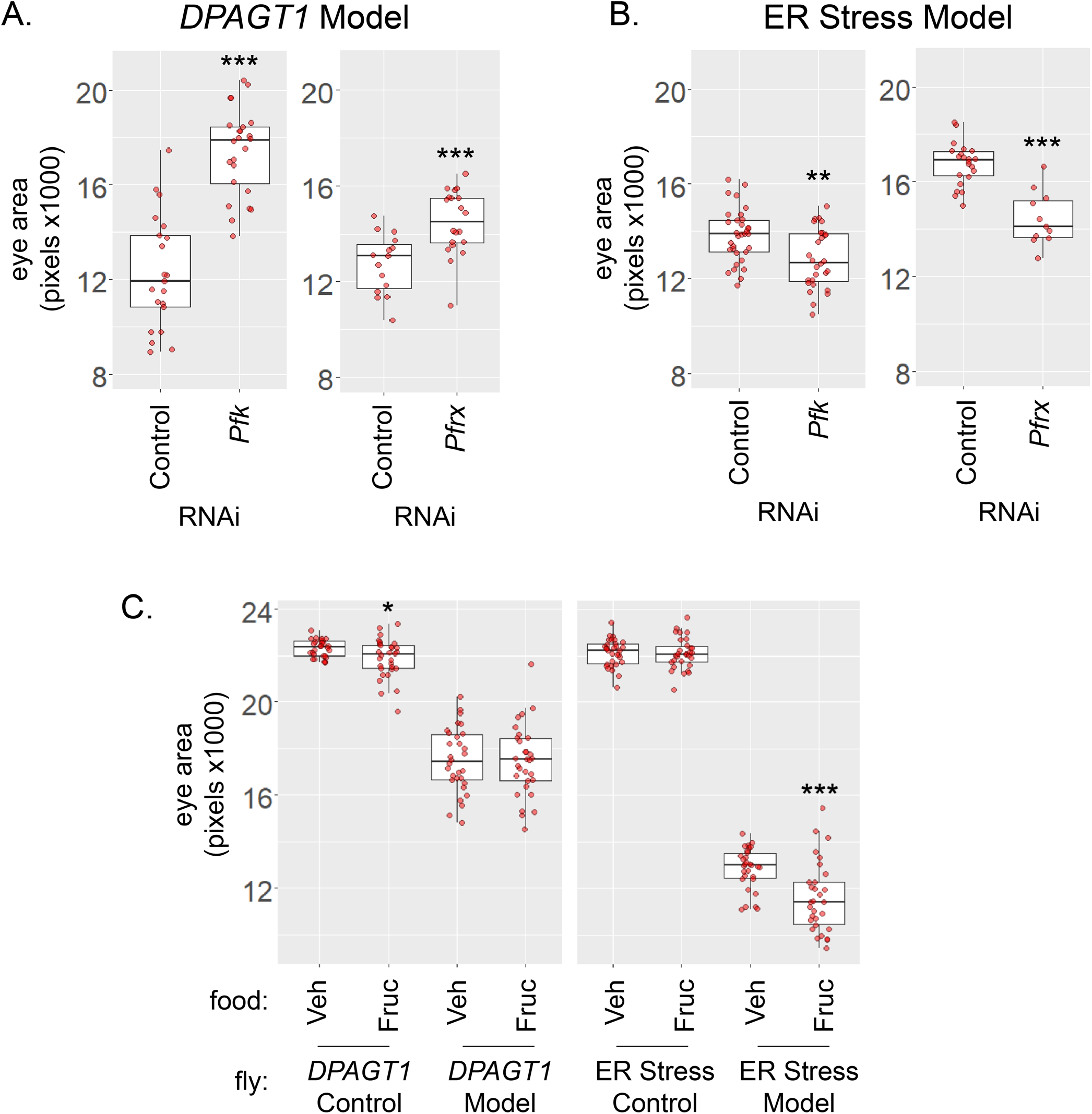
The source of ER stress can alter fructose metabolism. A-B. In the *DPAGT1* model, knockdown of *PFKM*/*Pfk* (BDSC 34336) and *PFKFB1*/*Pfrx* (BDSC 57222) causes a rescue of the *DPAGT1* deficiency, whereas in the ER stress model, knockdown of these genes causes a worse overall phenotype of eye size. C. Compared to its background control, feeding of 100mM fructose had a no deleterious effect on the *DPAGT1* model. However, 100mM fructose feeding significantly reduced eye size in the ER stress model. For A-C, 1 replicate is shown of 2 biological replicates (Supp. Table 9), * p< 0.05, ** p<0.01, *** p<0.001 (Student’s t test).

As previously discussed, knockdown of either of the two phosphofructokinases, *Pfk* and *Pfrx*, improved the *DPAGT1* model (Fig 5A, Supp. Table 7). Remarkably, however, knockdown of either of these two phosphofructokinases worsened the ER stress model (Fig 5B, Supp. Table 7). Pfk and Pfrx both act on the early glycolytic metabolite fructose-6-phosphate as a substrate to regulate glycolysis (Fig. 1C). We expect that loss of *Pfk* and *Pfrx* function should increase the pool of fructose in the cell. Given the shared substrate and differential effects in the two models, there may be key differences in how increased fructose levels and altered metabolism affects the loss of *DPAGT1* function and ER stress.

To test whether fructose differentially affects the loss of *DPAGT1* function and ER stress, we tested whether dietary supplementation of the two *in vivo* models with additional fructose might result in differential outcomes. In agreement with the RNAi data, feeding the ER stress model flies dietary fructose led to a significant decrease in overall eye size (−9.7% averaged from two biological replicates) (Fig. 5C, Supp. Table 9). In contrast, the genetically-matched control of the ER stress model had no effect from being fed dietary fructose. On the other hand, fructose supplementation resulted in no change in *DPAGT1* model eye size (Fig. 5C, Supp. Table 9). There was a small, but significantly negative, effect in the control-matched genetic background (−2.4%), suggesting that the lack of a difference in the *DPAGT1* model is in fact a net positive effect. The RNAi and fructose supplementation data together reveal a striking difference in fructose metabolism between the two models.

**Supplementary Figure 1. Qualitative comparisons of primary GDP-mannose, hexosamine, and glycolysis RNAi lines in *DPAGT1* and ER stress models**.

**Supplementary Figure 2. Qualitative comparisons of primary GDP-mannose, hexosamine, and glycolysis RNAi lines in *DPAGT1* and ER stress model control lines**.

**Supplementary Figure 3. Qualitative comparisons of downstream *Dpm1* glycosylation pathway RNAi lines in *DPAGT1* and ER stress models**.

**Supplementary Figure 4. Qualitative comparisons of downstream *Dpm1* glycosylation pathway RNAi lines in *DPAGT1* and ER stress model control lines**.

**Supplementary Figure 5. Simplified model of glycolysis and TCA cycle with labels for gene knockouts causing resistance or sensitivity**.

**Supplementary Table 1. List of all gene Log-fold change (LFC) data used in analysis and Volcano plot**.

**Supplementary Table 2. Comparison of fly orthologs of human CDG genes to survival screen candidate genes**.

**Supplementary Table 3. Gene Ontology (GO) analysis of all survival screen candidate genes**.

**Supplementary Table 4. List of all candidate genes from the ConA screen**.

**Supplementary Table 5. Gene Ontology (GO) analysis of all ConA screen candidate genes**.

**Supplementary Table 6. Comparison of candidate genes between the survival and ConA screens**.

**Supplementary Table 7. List of all primary candidate gene *in vivo* RNAi results**.

**Supplementary Table 8. Preliminary analysis of initial candidate genes *in vivo* RNAi lines**.

**Supplementary Table 9. Data on fructose feeding assay**.

## Discussion

Here, we report hundreds of new candidate genes that modify the effects of *DPAGT1* inhibition identified using a cell-based CRISPR screening approach. Testing these candidate genes *in vivo* validated many of the strongest hits and indicated that knockdown of GPI anchor biosynthesis and downstream *DPM1*/*Dpm1* glycosylation pathways can rescue the effects of *DPAGT1* deficiency and ER stress. Even though single gene mutations in these pathways are associated with specific CDGs, knockdown of many of these genes paradoxically improved negative phenotypes associated with *DPAGT1* inhibition and ER stress both *in vitro* and *in vivo*.

One hypothesis for this outcome is that impairing glycosylation pathways might slow down the overall synthesis of glycoproteins, thereby allowing cells to better withstand *DPAGT1* inhibition. Similar mechanisms are utilized in the response to ER stress, where the UPR will reduce protein synthesis in response to ER stress, giving the cell time to recover^3,43^. Another mechanism by which impairing glycosylation may rescue the loss of *DPAGT1* function is by freeing up cellular machinery that helps fold or degrade misfolded proteins. One such pathway is the calnexin/calreticulin (CNX/CRT) cycle, a process that ensures the proper folding of N-glycosylated proteins in the ER^44^. Recently, the CNX/CRT cycle was found to help mature GPI-associated proteins as well^45^. In one scenario, it is possible that impairing the GPI anchor biosynthesis pathway helps free up machinery in the CNX/CRT cycle, allowing it to fix more misfolded N-glycosylated proteins under loss of *DPAGT1* function. This might provide a novel approach to DPAGT1-CDG or other glycosylation disorders, i.e., a therapy might be designed to slow down global glycosylation and/or globally activating proteostasis machinery, to allow impacted cells more time to repair or clear misfolded proteins.

Because perturbation of each downstream Dpm1 glycosylation pathway alone was capable of partially rescuing the *DPAGT1* model, each one could provide a good therapeutic target for DPAGT1-CDG. Although targeting *DPM1* itself might be a useful approach, targeting a downstream gene may be better in practice to reduce the effects of elimination of all of its glycosylation pathways. For example, knockdown of *POMT2/tw* almost reached the same level of rescue as *Dpm1* alone (+45.9% vs. +62.1%), while targeting *POMT2* would perturb only O-mannosylation. O-mannosylation affects cadherins, plexins, and alpha dystrophin, among other proteins^46^, and is associated with multiple CDGs, including POMT2-CDG^47^. Thus, like many other candidate genes identified in our study, care would need to be taken in inhibiting this gene in any potential therapies. Nevertheless, given the multiple target genes throughout these pathways, this approach should allow for better “titration” of genetic or pharmacologic downregulation, which could hopefully provide an effective therapeutic approach.

RNAi against many of the GPI anchor biosynthesis genes improved one or both of the *in vivo* models. However, despite being a hit in both cell screens, *in vivo* knockdown of *PIGA/PIG-A* caused a reduction in eye size in the *DPAGT1* model and no change in the ER stress model. We hypothesize that *PIGA/PIG-A* may be too critical for development *in vivo*, such that developmental effects mask potential protection in either model. In support of this hypothesis, *PIGA/PIG-A* knockdown alone results in more severe eye phenotypes compared to other *PIG* genes (Supp. Fig. 3-4). There may simply be a limit on how strongly these downstream pathways can be inhibited before effects change from beneficial to deleterious.

The strongest candidate gene providing resistance to DPAGT1 inhibition was *PEK* (*PERK*). *PEK* encodes a protein kinase essential to the cellular ER stress response; it activates the transcription factor Crc (ATF4), which can regulate both pro-survival or apoptotic-related genes^5^. One hypothesis for why knockout of *PEK* leads to increased survival is that ER-stress-induced apoptotic-related gene expression might be lower in these cells, which may allow them time to grow and repair despite being stressed. While *Drosophila* lacks the ATF4 downstream pro-apoptotic target CHOP, *Drosophila* Crc can activate apoptosis by downregulation of the anti-apoptotic E3 ubiquitin ligase XIAP/Diap1^48^. Similar to *PERK*, knockout of *crc* does provide resistance (lfc = 0.75); however, knockout of its downstream target *XIAP/Diap1* surprisingly also caused resistance (lfc = 1.7), suggesting that this pathway is not the route of *PERK* knockout resistance. Interestingly, Crc is also a co-activator of the hormone receptor FXR/EcR^49^, and signaling of the hormone ecdysone through this receptor is critical for proper morphogenesis in the fly^50,51^. Knockout of *FXR/EcR* also provides resistance (lfc = 2.9). Thus, part of the resistance from *PEK* and *crc* knockout might result from its connection to ecdysone signaling.

There are several other candidate genes for which study of their roles in DPAGT1 deficiency might be informative. For example, another enriched GO pathway from the ConA screen was “inositol metabolic process” (GO:0006020) (Table 3). Inositol has several functions, including growth, immunity, and GTPase signaling^52^. One of these functions may play an important role in DPAGT1 inhibition, unrelated to glycosylation. Other GO enriched categories of note include nucleotide synthesis (Table 3). Activation of the UPR transcription factor ATF4 can promote *de novo* purine synthesis through the tetrahydrofolate cycle^53,54^. Perhaps modulation of purine synthesis might provide some benefit under DPAGT1 inhibition.

We also found that altering fructose metabolism had a different effect on the *DPAGT1* and ER stress models, and this might be explained by the mechanism of action in each model. The *DPAGT1* model ostensibly works by reducing the amount of functional DPAGT1 enzyme (similar to Tun treatment), and DPAGT1 enzymatic activity uses UDP-GlcNAc, which is a product of fructose metabolism through the hexosamine pathway (Fig.1C). Inhibition of DPAGT1 enzymatic action may result in a complex pattern of feedback that affects both fructose levels and metabolism. In contrast, the ER stress model overexpresses a single misfolded protein, causing accumulation in the ER lumen and subsequent ER stress^34^. The ER stress model does not directly impair the fructose metabolism pathway and its misfolded protein landscape is likely dominated by a single misfolded protein. Compared to the *DPAGT1* model, the ER stress model has a more ‘direct’ route to inducing ER stress. Given these differences, we hypothesize that altering fructose metabolism has a differential effect on DPAGT1 because of its metabolic connection to the hexosamine pathway.

In this study, we used cell culture and *in vivo* approaches to identify hundreds of new modifier genes affecting *DPAGT1* inhibition. Strikingly, we found that Dpm1 and its downstream glycosylation pathways are major enriched sites of these modifier genes. Our screen highlights that genes individually associated with one CDG may also be a source of modifiers of another CDG. If found to be more generally true, this paradoxical relationship between CDG genes could be exploited for therapeutic benefit. Targeting individual parts of these pathways under careful titration with drug or gene therapy may be the answer to better treatments for DPAGT1-CDG. In addition, we found differences in fructose metabolism that may explain differences between models of disease and ER stress and could be key to developing future dietary treatments for this disorder. Taken together, we believe these findings serve as a staging ground for further elucidation of modifier genes of *DPAGT1* inhibition and how they affect its underlying metabolism.

## Materials and Methods

### Cell culture

*Drosophila melanogaster* S2R+ cells are from by the Drosophila RNAi Screening Center (Harvard Medical School). A subline expressing SpCas9 and containing an attP integration site, S2R+/NPT005/MT-Cas9 (PT5/Cas9), was described previously^20,55^ and is available at the Drosophila Genomics Resource Center (Cell line # 268). Cells were maintained in Schneider’s media (Thermo 21720) supplemented with 1X Penn/Strep (Thermo 15070063) and 10% FBS (Thermo 16140071).

### Genome-wide CRISPR–Cas9 screening for resistance to tunicamycin

The *Drosophila* CRISPR genome-wide knockout library was previously described and is available from Addgene (134582-4)^20,55^. In brief, PT5/Cas9 cells were transfected with an equal-parts mixture of pLib6.4 containing an sgRNA library as well as pBS130 (26290, Addgene) using Effetene (301427, Qiagen) following the manufacturer’s instructions. The library was delivered in three parts, sublibrary group 1, 7,956 gRNAs for 995 genes; sublibrary group 2, 17,827 gRNAs for 2979 genes; and sublibrary group 3, 59,406 gRNAs for 9954 genes. After 4 days, the transfected cell library was selected with 5 μg/mL puromycin (540411, Calbiochem) for 12 additional days, subculturing every 4 days. After the stable sgRNA*-*expressing cell library was established, 1000 cells per sgRNA were subcultured in each passage to maintain the expected diversity of the sgRNA library. The cells were exposed to 590 nM tunicamycin (Cayman Chemical # 11445) for 30 days, passaging the same number of cells (∼8×10^7^) to four new 15 cm dishes every 4 days. Following the final passage, aliquots of the cells were collected, and their genomic DNA was extracted using a Zymo Quick-gDNA Miniprep kit (D3025, Zymo Research). DNA fragments containing the sgRNA sequences were amplified by PCR using at least 1000 genomes per sgRNA as template for each sample. The in-line barcoding strategy and sequences were as previously described^55^. Next generation sequencing (Illumina NextSeq) was performed at the Biopolymers Facility at Harvard Medical School. Following barcode demultiplexing, a readcount table was prepared from the raw sequencing files using MAGeCK version 0.5.9.4^56^, *count* subprogram, using standard parameters with 22 bp 5’ trimming and median-normalizing each sublibrary to 10,000 reads. The median log2 fold-change reported throughout was calculated from this data directly by measuring log2 of treated readcount divided by untreated readcount and then determining the median value among all sgRNAs targeting the same gene. For the volcano plot, the log2 fold-change and robust rank aggregation (RRA) p-value measures were determined using MAGeCK version 0.5.9.4 software, *test* subprogram, using standard parameters.

### FACS-based selection for altered concanavalin A staining intensity

After generating puromycin-resistant pools of cells expressing a genome-wide library CRISPR sgRNA library (88,627 sgRNAs targeting 13,685 *Drosophila* genes), the cells were again subjected to treatment with tunicamycin (590 nM) in normal growth medium for 1 week. Next, 1×10^8^ cells were placed into suspension in 10 mL and labeled live with 10 μg/mL Alexa Fluor 647 Conjugated Concanavalin A (Thermo # C21421) for at least 2 hours by diluting the stock 1:500 into full growth medium at room temperature with occasional inversion. 1 mL aliquots of labeled cells were transferred individually to 5 mL 40 μm filter-cap vials to mechanically declump the cells. Cells were then subjected to fluorescence-activated cell sorting (FACS) using a Sony MA900 FACS machine guided by gates established by tunicamycin-untreated cells stained in parallel and unstained cells. Approximately 1% of cells that exhibited the greatest intensity of staining in the AlexaFluor 647-A filter (corresponding to the average intensity of tunicamycin-untreated cells) were sorted into full growth media. Gating was reestablished after each 1 mL to account for increased uptake of the dye over the course of the experiment (∼6 hours). Finally, collected cells were transferred to a 6-well dish well and gently spun down at 100 x g for 10 min and their genomic DNA was extracted using the Zymo Quick-gDNA Miniprep kit, then subjected to PCR and Illumina sequencing as described. Amplicon sequencing was carried out at the MGH DNA core. A readcount table was prepared from raw sequencing files using MAGeCK version 0.5.9.4, *count* subprogram, using standard parameters with 22 bp 5’ trimming. Readcounts were sorted to reveal genes that had 4 or more sgRNAs in the top 1000 of all sgRNAs.

### Fly stocks and maintenance

All flies in this study were maintained at room temperature and fed a standard fly diet based on the Bloomington Drosophila Stock Center Standard Cornmeal Medium with malt (unless otherwise noted). Stocks obtained from the Bloomington Drosophila Stock Center were used in this study (listed in Supp. Table 7). The w-;; *eya composite*-GAL4 line was a gift from Justin Kumar (Indiana University Bloomington) and was characterized previously^57^. The “ER stress model” contains *GMR-GAL4* and *UAS-Rh1*^*G69D*^ on the second chromosome and has been previously described^34–36^.

The *DPAGT1* model was generated as follows. The *eya composite-*GAL4 and UAS-*Alg7* RNAi line (BDSC #53264) were crossed to create an *eya composite-*GAL4/UAS-*Alg7* RNAi (III) line. This line was then crossed to a balancer +/TM3, Dfd-YFP, *Sb* (III) and progeny were examined for crossover events. Progeny with degenerate eyes and *Sb* were collected, maintained for stability of the phenotype, and referred to as the *DPAGT1* model (*w-, y-, v-, sc-, sev-; +/+*; (*eya composite*-GAL4, w+, P[sc+, y+, v+, *Alg7* RNAi])/TM3, Dfd-YFP, *Sb*).

### Eye imaging and quantification

Adult female flies (2-7 days old) were collected under CO_2_ anesthesia, then placed on ice and transferred to − 80°C for later imaging. Eyes were imaged at 3x magnification (Leica EC3 Camera). Eye area was determined as previously described^35^. All measurements were done blinded to the RNAi line used, and one replicate from each model was measured by a second observer (99.2% agreement on average from 23 crosses, data not shown). Qualitative images of all control flies are also included as a reference (Supp. Fig. 2, 4). Note that when two RNAi lines were assayed together, the same control was compared to each. When possible, we used two different RNAi lines to limit the possibility of reagent-specific effects. Complete information on lines used can be found in Supp. Table 7 and representative images can be found in Supp. Fig. 1-4. In circumstances where only one RNAi line was available, when possible, multiple genes from the same pathway were tested in order to draw conclusions on the pathway as a whole in order to further limit any reagent-specific effects. For *Dpm1*, the BDSC 50713 RNAi line was lethal in the *DPAGT1* model and its control (Supp Table 7), and studies with the *Dpm1* BDSC 51396 RNAi line were performed at 18°C in the w-;; *eya* composite-GAL4 control line to increase viability. Both *Dpm1* RNAi lines were viable when crossed with the ER stress model.

### Fructose supplementation

To make the fructose-supplemented media, our standard media was melted in a microwave and maintained at 95°C on a stir plate. We used this melted media as-is for the control and added fructose to a final concentration of 100mM to a separate set of vials and stirred until it was fully dissolved. Flies were mated in these vials and removed after egg laying, allowing progeny to feed on sugar-supplemented media from hatching. Timing and quantitative imaging of these progeny was performed as described above.

### Statistics

Genetic overlap representation factors and probability statistics were calculated via “Nematode bioinformatics” (http://nemates.org). For the CDG gene overlap analysis, we used the DIOPT ortholog finder (https://www.flyrnai.org/cgi-bin/DRSC_orthologs.pl)^21^ and analyzed any hits with a DIOPT score of 5 or greater.

Gene Ontology (GO) analysis was performed with the PANTHER Overrepresentation test using the “GO biological process complete” data set, compared to all *Drosophila* genes, and using the default parameters (Fisher’s Exact, False Discovery Rate)^58,59^.

Eye size comparisons were analyzed using the Student’s t test. When comparing groups of RNAi lines to a single control, Bonferroni multiple testing correction was used. Venn diagram and graphs of eye size comparisons were made using the statistical software R^60^.

## Funding

This work was supported by NIGMS R35 GM124780 to CYC and P41 GM132087 to NP, and by a grant from the Primary Children’s Hospital Center for Personalized Medicine to CYC. HMD was supported by an NIGMS NRSA award (F32 GM136057). JSZ was supported by the Undergraduate Research Opportunities Program (UROP) through the University of Utah Office of Undergraduate Research. NP is an investigator of the Howard Hughes Medical Institute. The BDSC is supported by NIH P40OD018537.

## Acknowledgements

We thank Emily Coelho, Shayna Scott, and Bekah Rushforth for technical assistance with fly management.

## Notes

### Competing Interest Statement

The authors have declared no competing interest.

