## Supplementary figures and images for "A genome-wide CRISPR screen identifies the glycosylation enzyme DPM1 as a modifier of DPAGT1 deficiency and ER stress"

### Supplemental Figure 1

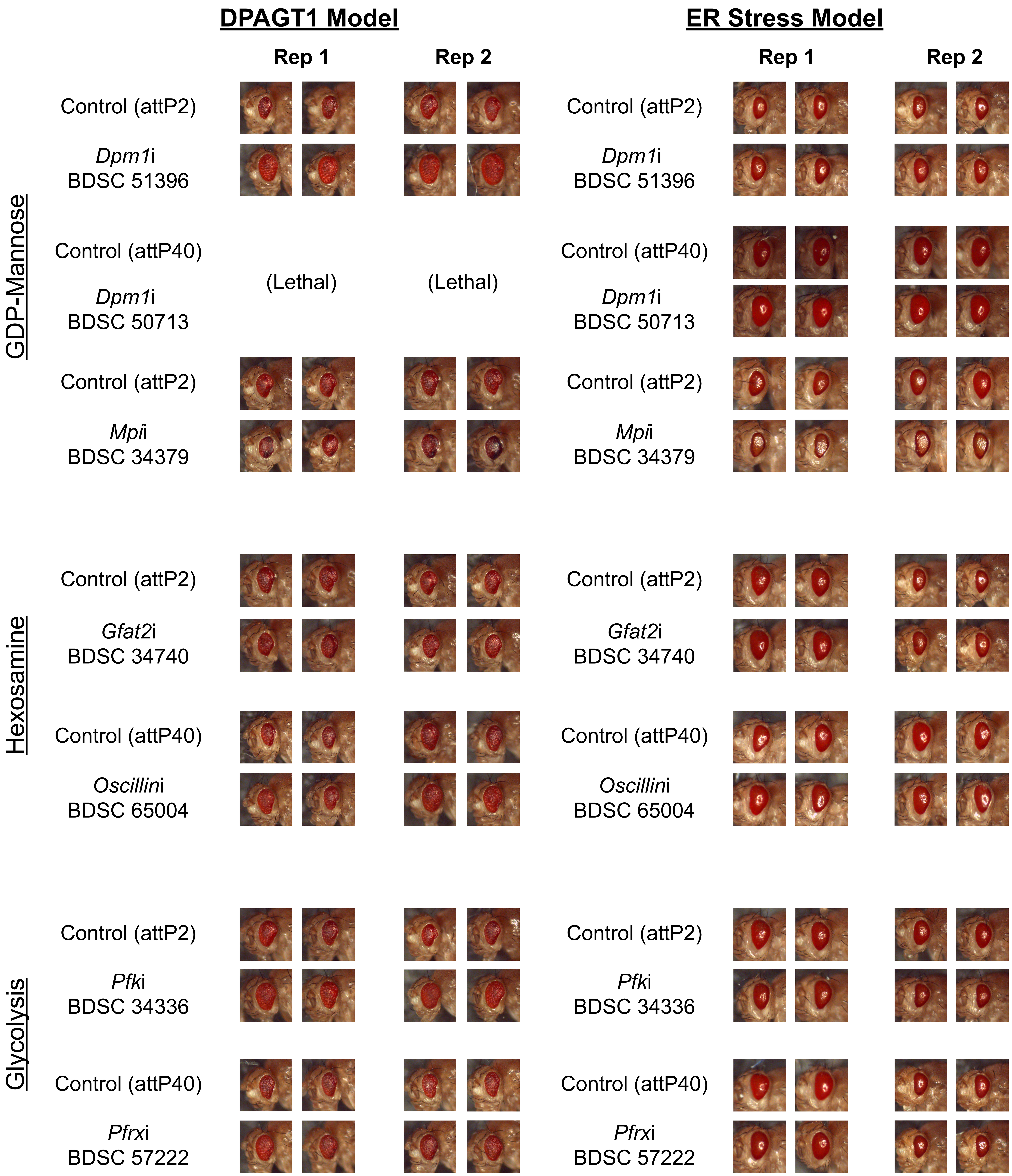

### Supplemental Figure 2

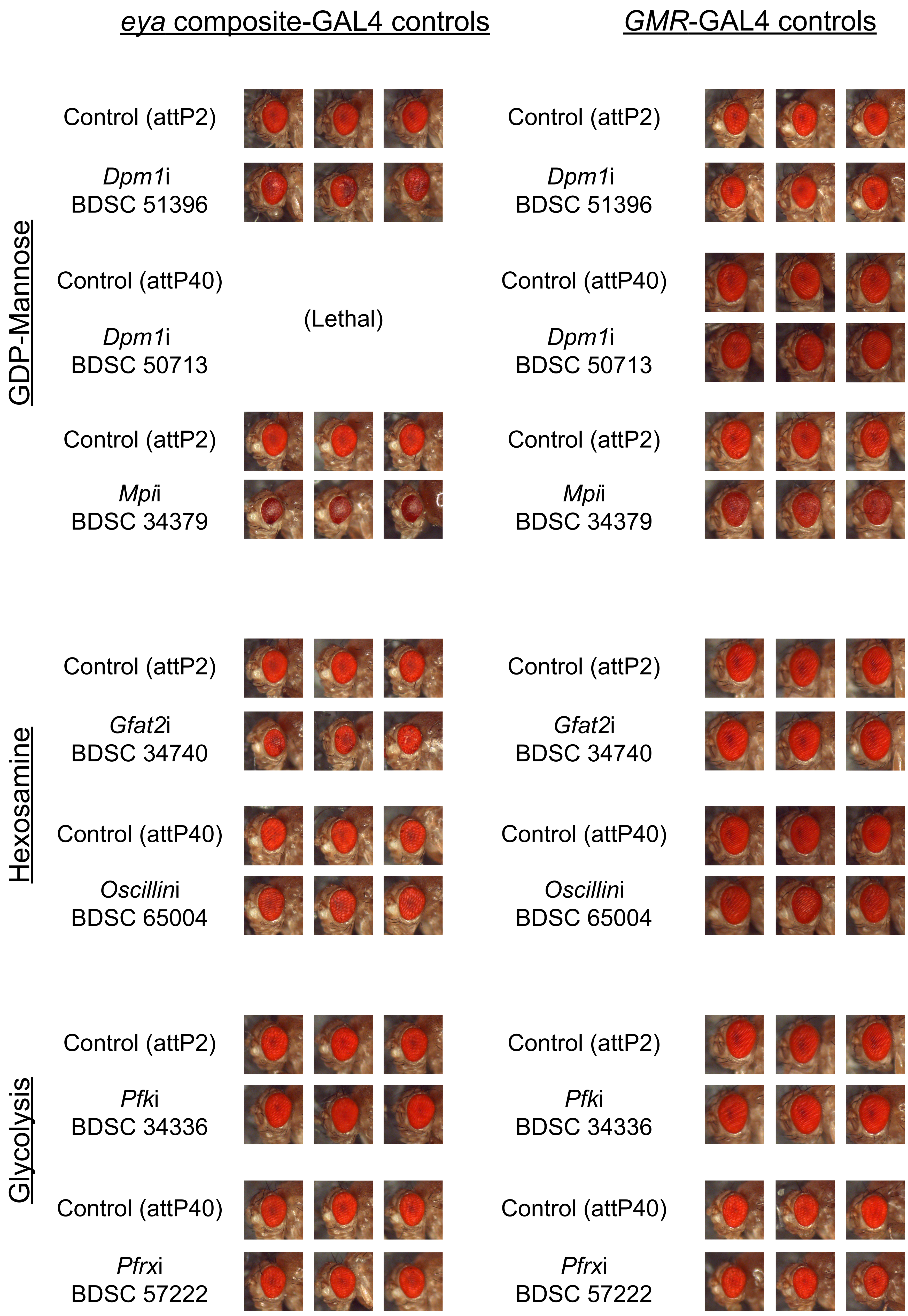

### Supplemental Figure 3

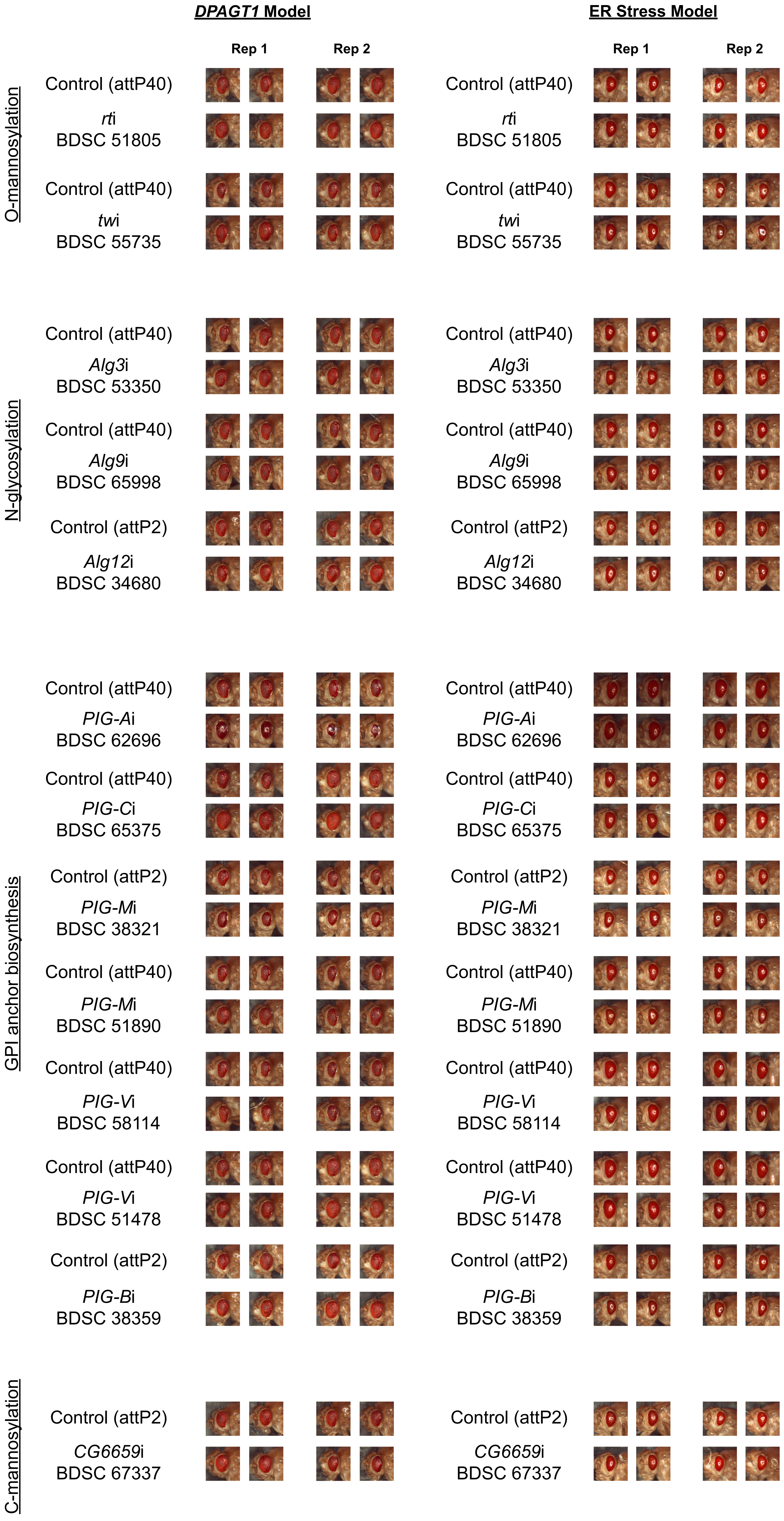

### Supplemental Figure 4

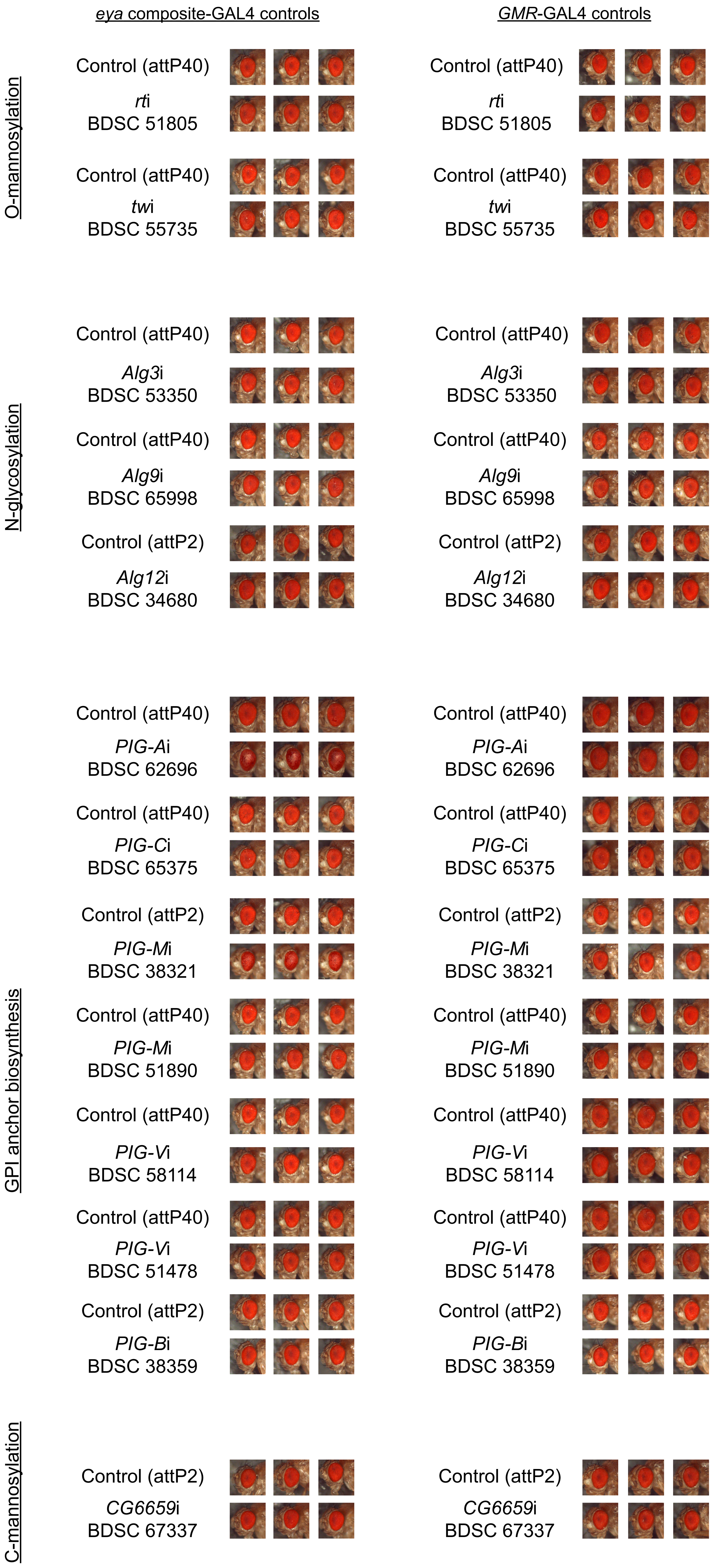

### Supplemental Figure 5

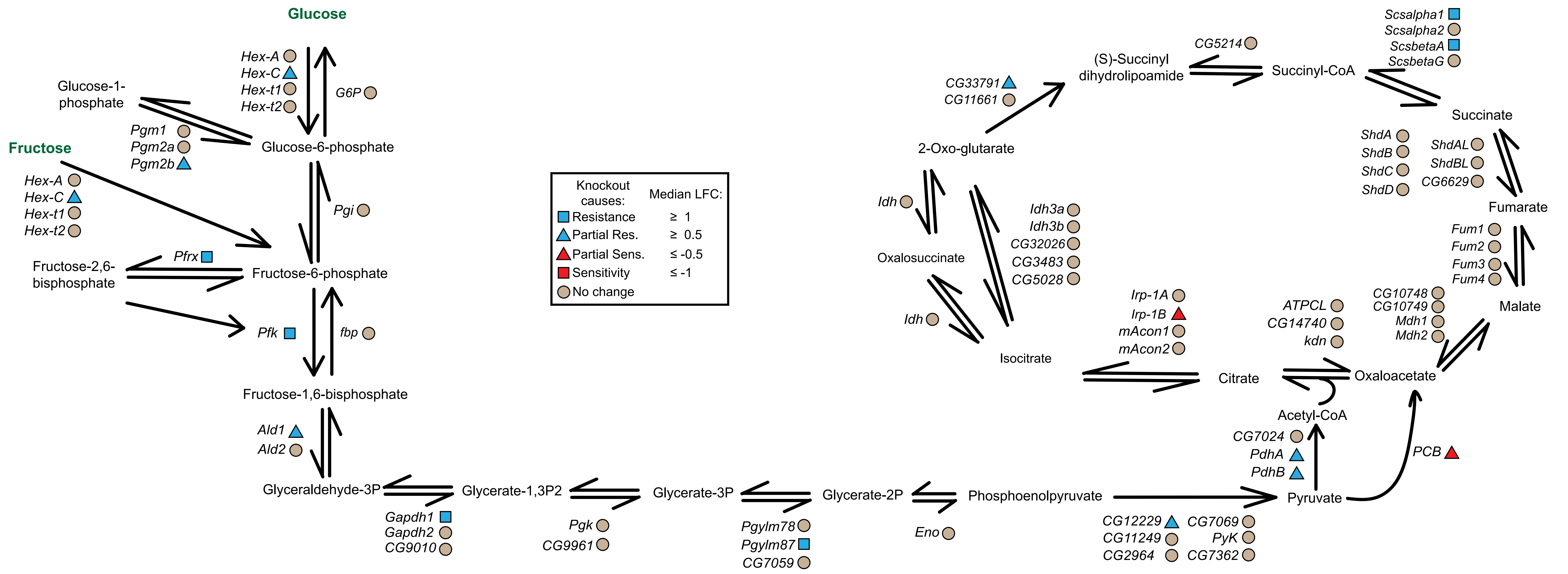
